# Disparities in spatially variable gene calling highlight the need for benchmarking spatial transcriptomics methods

**DOI:** 10.1101/2022.10.31.514623

**Authors:** Natalie Charitakis, Agus Salim, Adam T. Piers, Kevin I. Watt, Enzo R. Porrello, David A. Elliott, Mirana Ramialison

## Abstract

Identifying spatially variable genes (SVGs) is a key step in the analysis of spatially resolved transcriptomics (SRT) data. SVGs provide biological insights by defining transcriptomic differences within tissues, which was previously unachievable using RNA-sequencing technologies. However, the increasing number of published tools designed to define SVG sets currently lack benchmarking methods to accurately assess performance. This study compares results of 6 purpose-built packages for SVG identification across 9 public and 5 simulated datasets and highlights discrepancies between results. Additional tools for generation of simulated data and development of benchmarking methods are required to improve methods for identifying SVGs.

## Background

Spatially resolved transcriptomics (SRT) captures variations in gene expression across tissues (1–7). Computational methods for analysis of SRT data are being established (8–12). Among the goals of SRT analysis pipelines is the robust and reliable identification of spatially variable genes (SVGs) within tissue sections (12–14). SVGs are defined as having expression levels across a tissue that covary in a location specific manner (13,15). Published methods for SVG identification employ different mathematical models aiming to capture biological truth (13,14,16–27). Benchmarking of analysis tools is needed to ensure the reliability of processed data matches or supersedes that of similar technologies such as single-cell (sc) RNA-Seq (28,29).

Here, we compared the performance of six packages – SpatialDE (13), SPARK-X (16), Squidpy (30), Seurat (19), SpaGCN (18), scGCO (31) from healthy and cancerous fresh frozen (FF) and formalin-fixed paraffin embedded (FFPE) tissues generated with 10X Visium technology alongside simulated datasets (Additional file 1) (32,33). Our analysis reveals discrepancies in SVGs identified by each package highlighting the urgent need for further benchmarking and development of tools to identify SVGs.

## Results and Discussion

Comparing SVGs predicted by six packages, we found that the numbers of SVGs identified differed by one to two orders of magnitude across the same tissue for all nine investigated datasets (Fig. 1A & Additional files 2-4). For example, in the FF endometrial adenocarcinoma ovarian tissue dataset, the number of predicted SVGs ranged from 87 to 3707 (Fig. 1A). For 7/9 datasets, SpatialDE identifies the greatest number of SVGs, while SpaGCN identifies the fewest number of SVGs in 6/9 public datasets (Fig. 1A & Additional files 2-4). Apart from the FF mouse brain coronal section dataset, packages tested tend to report more SVGs in cancerous datasets than in healthy tissues, most evident across SpatialDE and SPARK-X (Additional file 5). There is no apparent difference in reported SVGs driven by data generated from FF or FFPE tissue (Additional file 5).

**Figure 1.**
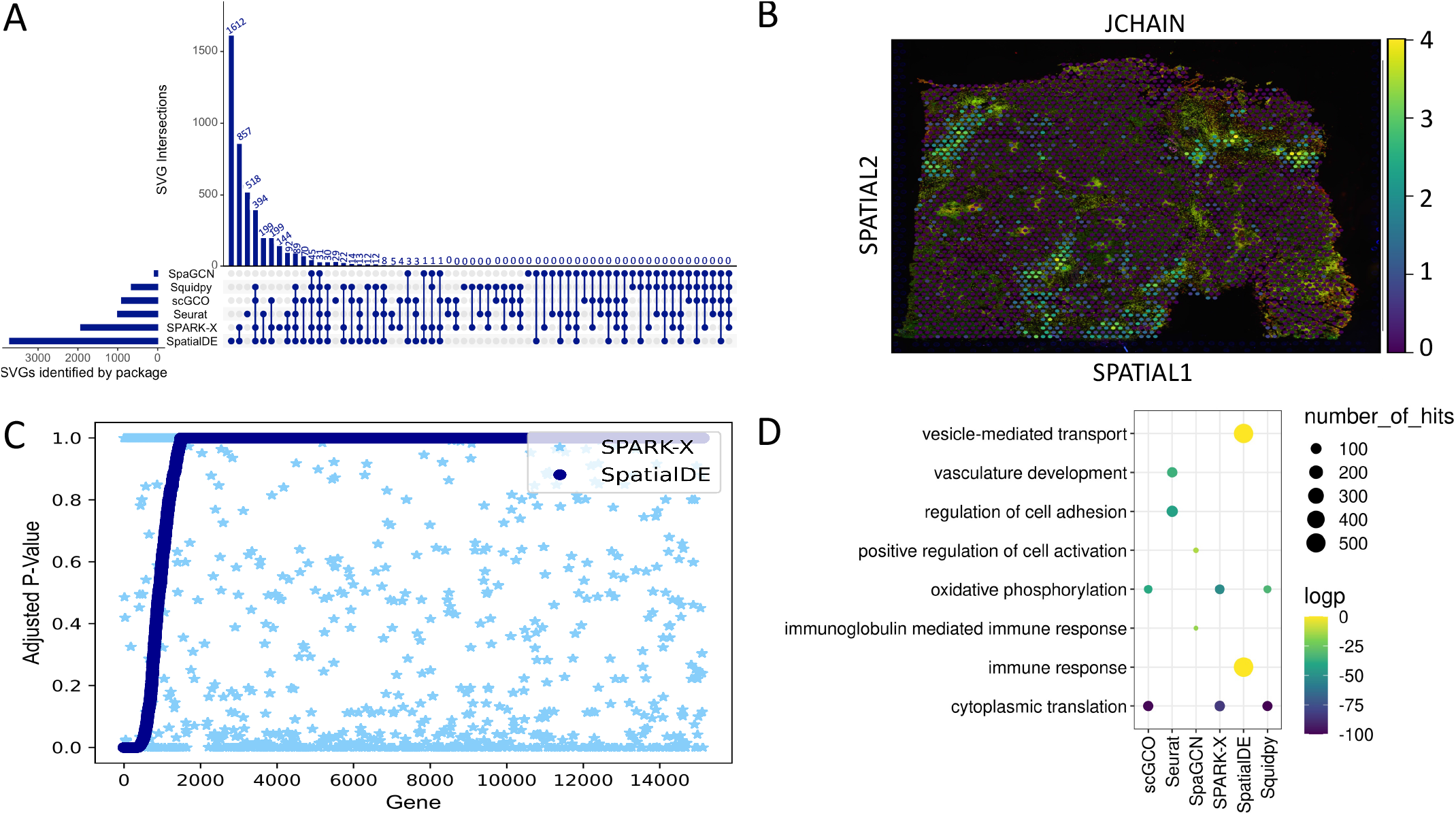
Discrepancies between SVGs in a dataset annotated by six different packages. **A)** Upset plots displaying the distinct intersections of SVGs identified in FF endometrial adenocarcinoma ovarian tissue dataset when analysed with SpaGCN, Squidpy, scGCO, Seurat, SPARK-X and SpatialDE. **B)** Pattern of expression of *JCHAIN*, a SVG identified by all six packages in FF endometrial adenocarcinoma ovarian tissue. Spots representing the capture regions of the Visium slide are overlayed on accompanying histological image. **C)** Sorted q-values from SpatialDE results across the FF left ventricle datasets plotted against SPARK-X q-values for the same gene. **D)** Gene ontology enrichment results from FF endometrial adenocarcinoma ovarian tissue dataset using SVGs identified by each package as inputs.

**Figure 2.**
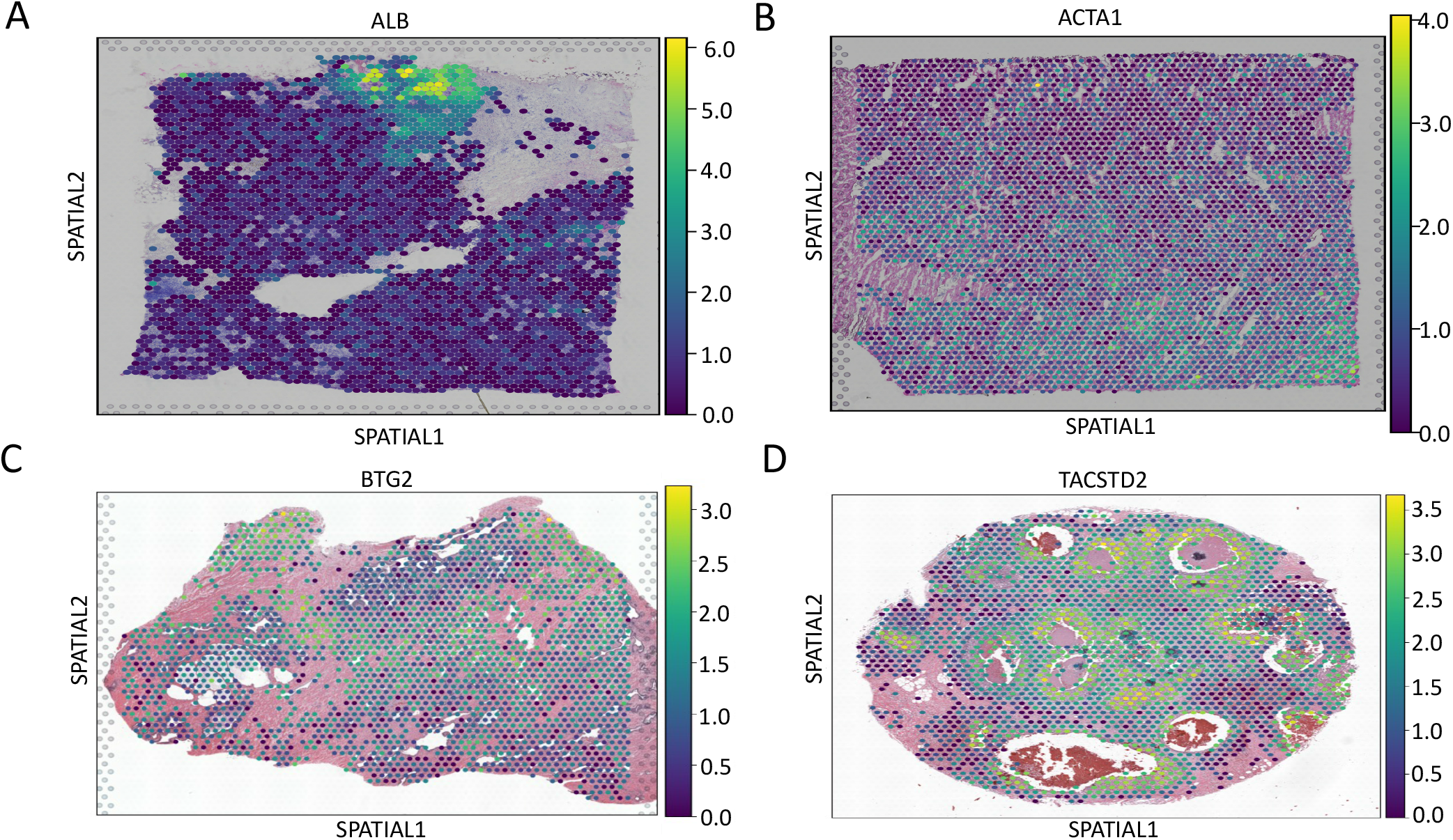
Expression patterns of SVGs identified by all tested packages in a single dataset. **A)** Pattern of expression of *ALB*, a SVG identified by all six packages in FF invasive ductal carcinoma breast tissue. **B)** Pattern of expression of *ACTA1*, a SVG identified by all six packages in FF left ventricle. **C)** Pattern of expression of *BTG2*, a SVG identified by all six packages in FFPE prostate. **D)** Pattern of expression of *TACSTD2*, a SVG identified by all six packages in FFPE invasive ductal carcinoma breast tissue.

To understand the disparity in the number of SVGs reported, we hypothesised that methods used to correct for type I error might influence results. SpatialDE, Squidpy, scGCO and SPARK-X report a q-value/FDR for all genes. Methods that do not supply associated p-values report the lowest number of SVGs in half of the datasets (Fig. 1A & Additional files 2-4). Across all datasets, a minimal number of common SVGs are identified between all 6 packages that each display an expression pattern that aligns with spatial correlation suggesting that they are *bona fide* SVGs (Fig. 1B).

Next, to assess the significance of small overlap between predicted SVGs, the ranked lists of SVGs provided by SpatialDE, Squidpy, scGCO and SPARK-X were compared across datasets using a Wilcoxon signed rank test. We observed differences in the SVG ranking across the tested packages, indicating that the lack of overlap is statistically significant (Fig. 1C & Additional files 6-11, Additional file 12: Table S1). q-values for a single gene differ between packages (Fig. 1C & Additional files 6-11).

We investigated the downstream impact arising from discordant results by comparing gene ontology (GO) enrichment analysis using SVGs predicted by each package within the same tissue. (Fig. 1D & Additional file 13). We observed non-overlapping parent terms between datasets in the top 10 over-represented GO terms (Fig. 1D & Additional file 13). These results suggest that gene sets predicted by each package do not overlap functionally. Further, restricting GO analysis to SVGs can bias downstream interpretation of results, especially if there is a risk of a large percentage of false positives (FP).

Despite the discrepancies, SVGs identified by all packages within the same dataset display an expression pattern aligned with a clear spatial correlation. An example is *JCHAIN*, which is expressed in endometrial macrophages and associated with adenocarcinoma (Fig. 1B) (34–36). Furthermore, in FF mouse brain coronal section, a dataset with known spatial gene expression patterns, many of the 368 commonly identified SVGs display expression patterns corresponding to their known region of expression in the Allen Mouse Brain Atlas (Additional file 13) (37). These observations indicate that patterns with a stronger signal-to-noise ratio are identified by all methods.

To gain insight into the inconsistent results between different packages we evaluated their sensitivity and specificity. To determine true positives and FP rates, we first generated two simulated datasets with different known SVG patterns with SRTsim (Additional file 15: A-B, Supplementary methods) (38). Each dataset contained 1500 known SVGs (Supplementary methods). Using each as the input to four packages (scGCO, SpatialDE, Squidpy and SPARK-X), all packages identify all true positives across datasets, except for Squidpy when analysing the dataset with lower signal SVGs. This indicates a potential need for a more relaxed Moran’s I statistic threshold (Additional file 12: Table S2). Additionally, most packages introduce minimal levels of FPs with simulated datasets, except for scGCO (Additional file 12: Table S2). This is possibly due to of the type I error correction method or sequential search procedure employed by scGCO (31).

To further investigate the potential of each package to call FPs, we generated four negative control datasets from two publicly available datasets. For each, two versions of negative controls were generated -[1] where spatial coordinates are permutated and [2] where spatial coordinates and columns of gene expression matrix are permutated. For all negative control datasets generated from the FF left ventricle, Seurat, scGCO, Squidpy and SPARK-X identified no SVGs, indicating an ability to distinguish signal from noise. SpaGCN identified <10 SVGs. SpatialDE identified 4 SVGs when both spatial coordinates and columns of the gene expression matrix are permutated and 2953 in the other dataset. For negative control datasets generated from FFPE prostate randomising only spatial coordinates, SpatialDE identified 632 SVGs, SpaGCN identified 232 and Squidpy identified 1. Seurat, scGCO and SPARK-X did not run with these inputs. When spatial coordinates and gene expression values were randomised, SpatialDE was the only package to identify SVGs (713). Patterns of the top SVGs identified by SpatialDE lack a strong spatial pattern (Additional file 16). This would indicate that SpatialDE is prone to introducing FPs into a dataset when genes are lowly expressed compared. However, the original SVGs identified by SpatialDE and the result in Fig. 1B suggest the top ranked SVGs labelled by SpatialDE in real data display a signal consistent with that of a SVG.

Inconsistencies in the results between different packages may therefore be explained by the differences in FP rates, rather than the rate of false negatives. To investigate this, expression of known housekeeping gene *Eef2* (39) was visualised across the FF mouse brain coronal Section dataset to verify its ubiquitous expression (Additional file 14: D). *Eef2* was found to be annotated as an SVG by scGCO, SPARK-X, SpatialDE and Squidpy. This indicates genes with high levels of expression, but that do not display strong spatial expression patterns, may be a source of confounding.

In addition to differences in FP rates, results generated from different packages applied to the same dataset may arise due to the different assumptions regarding the underlying distributions of gene expression and effectiveness of normalisation methods (Additional file 12: Table S3). Negative binomial distributions can successfully model sequencing-based methods that employ unique molecular identifiers (UMIs) and most gene expression patterns in Visium datasets (38,40). While certain packages implement their own normalisation method, there is no guarantee that data pre-processed using a separate workflow will have been normalised to fit the assumptions of gene distributions held by a particular package (13,19). Once methods for generating simulated SRT datasets with different underlying distributions of gene expression are published, the effect this has on identifying SVGs needs to be investigated thoroughly. There are many caveats to generating simulated datasets with known patterns of gene expression with efforts focused on simulating data with defined cell types (41–48).

## Conclusions

While different mathematical models for detecting SVGs produce varied results when applied to the same tissue, and across tissues and biological conditions, SVGs with a strong spatial covariance are consistently identified. This indicates that differences between methods do not bias performance in a particular tissue or biological condition. However, more transcriptionally heterogenous tissues appear to affect the performance of certain packages in calling SVGs and genes with unclear spatial patterns but high expression levels may be labelled as FP (14,15). Only four of the tested packages produced q-value ranked gene lists, thus differences in FP rates could confound comparisons of results across packages. Furthermore, due to the resolution of the Visium technology platform the presence of heterogenous cell types could complicate accurate identification of SVGs due to inherent transcriptional variation. Increased resolution may present challenges in terms of computational performance and the mathematical difficulties of dealing with highly sparse datasets. As SVG identification becomes an integral part of the computational analysis of SRT and datasets grow, a subset of SVGs could be used in preference over the entire dataset for downstream analysis (such as highly variable genes in scRNA-Seq analysis). Introducing noise may hamper key downstream steps in understanding spatially restricted disease states as well as novel targets for treatment (49,50). Methods that explore the entire dataset for SVG labelling paired with stricter q-value cut-offs (q-value < 0.01) can be employed when analysing tissues with known transcriptional complexity to decrease the possibility of FP being reported. Future work will include comparison of package performance across datasets generated from different SRT platforms. The development of workflows for simulating datasets and benchmarking current and novel methods for SVG identification will allow for accurate determination of the spatially restricted transcriptional differences that manifest as biological outcomes during development and disease.

## Methods

Six packages were selected for comparison: SpatialDE (13), SPARK-X (16), Squidpy (30), Seurat (19), SpaGCN (18) and scGCO (31), which each employ a different mathematical model to identify SVGs. Each package was tested on 9 publicly available, V1 Chemistry Visium datasets generated from healthy and cancerous tissues, most human and one mouse (Additional file: Table S4). The filtered output files from the Space Ranger v1.0.0 pipeline were downloaded, and pre-processed using Scanpy v1.8.1 (51). Two reference-free simulated datasets with features similar to Visium datasets were generated using SRTsim (38). Using the FF left ventricle and FFPE prostate datasets as inputs, two negative control datasets were generated in each instance by 1) independently randomising the x and y spot coordinates to remove spatial correlation and 2) additionally randomising each column in the gene expression matrix to ensure any overall association was removed.

## Declarations

Ethic approval and consent to participate

NA

Consent for publication

NA

Availability of data and materials

The input files for the Visium data used in this study are available from the <u>10X Genomics website</u>. Examples of the data pre-processing and analysis to identify SVGs on a single dataset for each of the six packages can be found on Github at the repository<u>Ramialison-Lab/Disparities_in_SVG_calling.</u>

Additional file 1: additional_file1.pdf

Overview of the design of benchmarking workflow. First, publicly available datasets are obtained to be used as inputs for pre-processing in a Scanpy workflow. For each dataset the processed output is then transformed into a data object most suitable for each of the SVG analysis packages. Simulated data is directly used as inputs for SVG analysis. Finally, the results are compared across packages within each dataset.

Additional file 2: additional_file2.pdf

Distinct overlap of SVGs identified by different combinations of the six tested packages. A) FF cerebellum. B) FF lymph node. C) FFPE adenocarcinoma prostate.

Additional file 3: additional_file3.pdf

Distinct overlap of SVGs identified by different combinations of the six tested packages. A) FF invasive ductal carcinoma breast tissue. B) FF left ventricle. C) FFPE prostate.

Additional file 4: additional_file4.pdf

Distinct overlap of SVGs identified by different combinations of the six tested packages. A) FFPE invasive ductal carcinoma breast tissue. B) FF mouse brain coronal section.

Additional file 5: additional_file5.pdf

Comparison of the number of SVGs identified by six different packages across different datasets. FFPE tissues are visualised by a cross and FF tissues are visualised with a square. High numbers of SVGs are identified in both FF and FFPE tissues, however tissues that are more transcriptionally complex seem to have more SVGs called across the dataset. Seurat reports a consistent number of SVGs across datasets as it first identifies highly variable genes then ranks their expression by how dependent it is on spatial location (52).

Additional file 6: additional_file6.pdf

Comparison of ranked SpatialDE q-values against gene-matched SPARK-X q-values of all genes generated from each dataset. Order of plots repeats across all datasets. A) FF cerebellum. FF lymph node. C) FFPE adenocarcinoma prostate. D) FF invasive ductal carcinoma breast tissue. E) FFPE prostate. F) FFPE invasive ductal carcinoma breast tissue. G) FF endometrial adenocarcinoma ovarian tissue. H) FF mouse brain coronal section.

Additional file 7: additional_file7.pdf

Comparison of ranked SpatialDE q-values against gene-matched scGCO q-values of all genes generated from each dataset. Order of plots repeats across all datasets. A) FF cerebellum. B) FF lymph node. C) FFPE adenocarcinoma prostate. D) FF invasive ductal carcinoma breast tissue. E) FFPE prostate. F) FFPE invasive ductal carcinoma breast tissue. G) FF endometrial adenocarcinoma ovarian tissue. H) FF left ventricle. I) FF mouse brain coronal section.

Additional file 8: additional_file8.pdf

Comparison of ranked SPARK-X q-values against gene-matched scGCO q-values of all genes generated from each dataset. Order of plots repeats across all datasets. A) FF cerebellum. B) FF lymph node. C) FFPE adenocarcinoma prostate. D) FF invasive ductal carcinoma breast tissue. E) FFPE prostate. F) FFPE invasive ductal carcinoma breast tissue. G) FF endometrial adenocarcinoma ovarian tissue. H) FF left ventricle. I) FF mouse brain coronal section.

Additional file 9: additional_file9.pdf

Comparison of ranked SpatialDE q-values against gene-matched Squidpy q-values of all genes generated from each dataset. Order of plots repeats across all datasets. A) FF cerebellum. B) FF lymph node. C) FFPE adenocarcinoma prostate. D) FF invasive ductal carcinoma breast tissue. E) FFPE prostate. F) FFPE invasive ductal carcinoma breast tissue. G) FF endometrial adenocarcinoma ovarian tissue. H) FF left ventricle. I) FF mouse brain coronal section.

Additional file 10: additional_file10.pdf

Comparison of ranked SPARK-X q-values against gene-matched Squidpy q-values of all genes generated from each dataset. Order of plots repeats across all datasets. A) FF cerebellum. B) FF lymph node. C) FFPE adenocarcinoma prostate. D) FF invasive ductal carcinoma breast tissue. E) FFPE prostate. F) FFPE invasive ductal carcinoma breast tissue. G) FF endometrial adenocarcinoma ovarian tissue. H) FF left ventricle. I) FF mouse brain coronal section.

Additional file 11: additional_file11.pdf

Comparison of ranked scGCO q-values against gene-matched Squidpy q-values of all genes generated from each dataset. Order of plots repeats across all datasets. A) FF cerebellum. B) FF lymph node. C) FFPE adenocarcinoma prostate. D) FF invasive ductal carcinoma breast tissue. E) FFPE prostate. F) FFPE invasive ductal carcinoma breast tissue. G) FF endometrial adenocarcinoma ovarian tissue. H) FF left ventricle. I) FF mouse brain coronal section.

Additional file 12: additional_file12.pdf

Contains Tables S1-4. Table S1) Reported statistic and associated p-value from each combination of pairwise comparison of results between SpatialDE, SPARK-X and scGCO when reported p-values were compared using the Wilcoxon signed rank test. Table S2) Sensitivity and specificity of scGCO, Squidpy, SPARK-X and SpatialDE when used to analyse both simulated datasets generated with SRTsim. Sensitivity and specificity are reported over a range of q-value cut-offs, from 0.01-1.0. Table S3) Packages included for identification of SVGs in benchmarking study. The packages are ordered by assumptions made on distribution of gene expression. The rows highlighted in blue indicate grouped packages using graph-based methods. Table S4) Overview of publicly available 10X Visium datasets to be included in benchmarking process. FF indicates tissues that are fresh frozen and FFPE indicates tissues that are formalin-fixed paraffin-embedded. The filtered output files and imaging data from the Space Ranger v1.0.0, v1.2.0 or V1.3.0 pipeline were downloaded for each dataset.

Additional file 13: additional_file13.pdf

Gene ontology enrichment results using SVGs identified by each package as inputs across datasets. A) FF cerebellum. B) FF lymph node. C) FFPE adenocarcinoma prostate. D) FF invasive ductal carcinoma breast tissue. E) FF left ventricle. F) FFPE prostate. G) FFPE invasive ductal carcinoma breast tissue. H) FF mouse brain coronal Section.

Additional file 14: additional_file14.pdf

Spatial expression patterns of SVGs identified by all packages across the FF mouse brain coronal section dataset. A) Expression of *Hap1* across the hypothalamus and amygdala, cross-referenced with the Allen Mouse Brain Reference. B) Expression of *Prkcd* localised to the thalamus, cross-referenced with the Allen Mouse Brain Reference. C) Expression of *Itpka*, with highest expression in the isocortex, hippocampal formation (HPF) and cortical subplate consistent with patterns displayed in the Allen Mouse Brain Reference. D) Expression of *Eef2*, a known housekeeping gene in mouse (39).

Additional file 15: additional_file15.pdf

Simulated datasets generated with SRT sim. A) Location of simulated SVGs with a hotspot pattern visualised in blue, while red area indicates expression of noise genes. B) Location of simulated SVGs in both blue and green corners, while red area indicates expression of noise genes. C) Distinct overlap of SVGs compared to the control SVG list identified by different combinations of the four tested packages. 1500 SVGs were present in this dataset. D) Distinct overlap of SVGs compared to the control SVG list identified by different combinations of the four tested packages. 750 high signal SVGs and 750 low signal SVGs were present in this dataset.

Additional file 16: additional_file16.pdf

Top SVG identified by SpatialDE in negative simulated datasets generated by A) Randomising coordinates of FF Left Ventricle dataset. B) Randomising counts and coordinates of FF left ventricle dataset. C) Randomising coordinates of FFPE prostate dataset. D) Randomising counts and coordinates of FFPE prostate dataset.

Additional file 17: additional_file17.pdf

Upset plot of results generated from running SPARK-X and SpatialDE on simulated data with known pattern of SVGs.

Additional file 18: additional_file18.xlsx Supplementary methods

## Authors’ contributions

N.C. performed experiments with contribution from A.S. on statistical design; N.C., A.S., A.T.P., K.I.W, E.R.P., D.A.E., M.R. designed experiments; N.C., K.I.W, D.A.E., M.R. wrote the manuscript with input from A.S. and E.R.P.

## Acknowledgements

The authors would like to thank Matthew Ritchie, David Powell and Gordon Smyth for their contributions of ideas to include in the analysis, Hieu Nim for manuscript feedback, Michael See for use of his code in generating plots and members of Transcriptomics and Bioinformatics and Heart Regeneration and Disease laboratories for their feedback.

