## Supplementary figures and images for "Disparities in spatially variable gene calling highlight the need for benchmarking spatial transcriptomics methods"

### Additional File 1

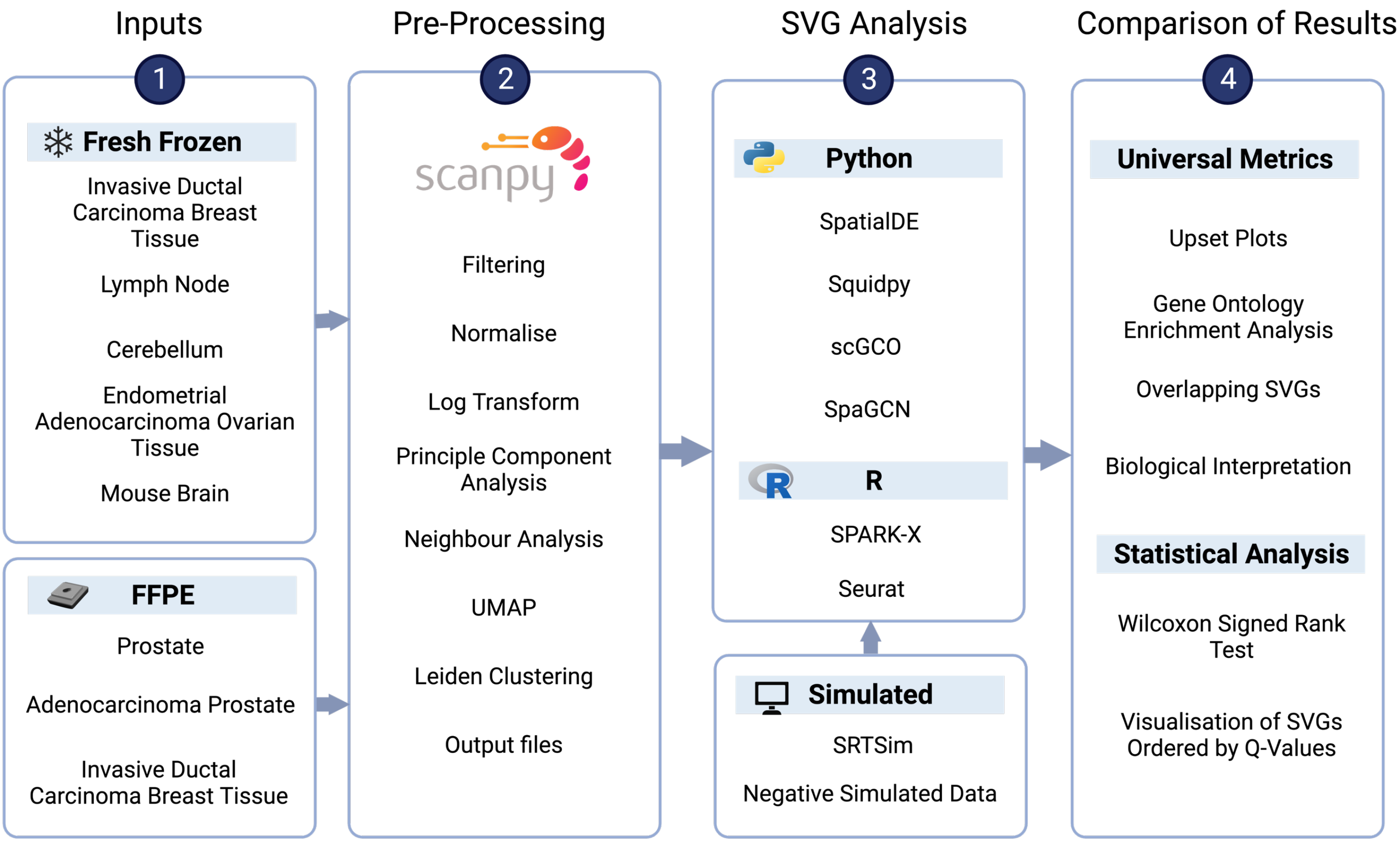


Charitakis et al. Fig. S1

### Additional File 2

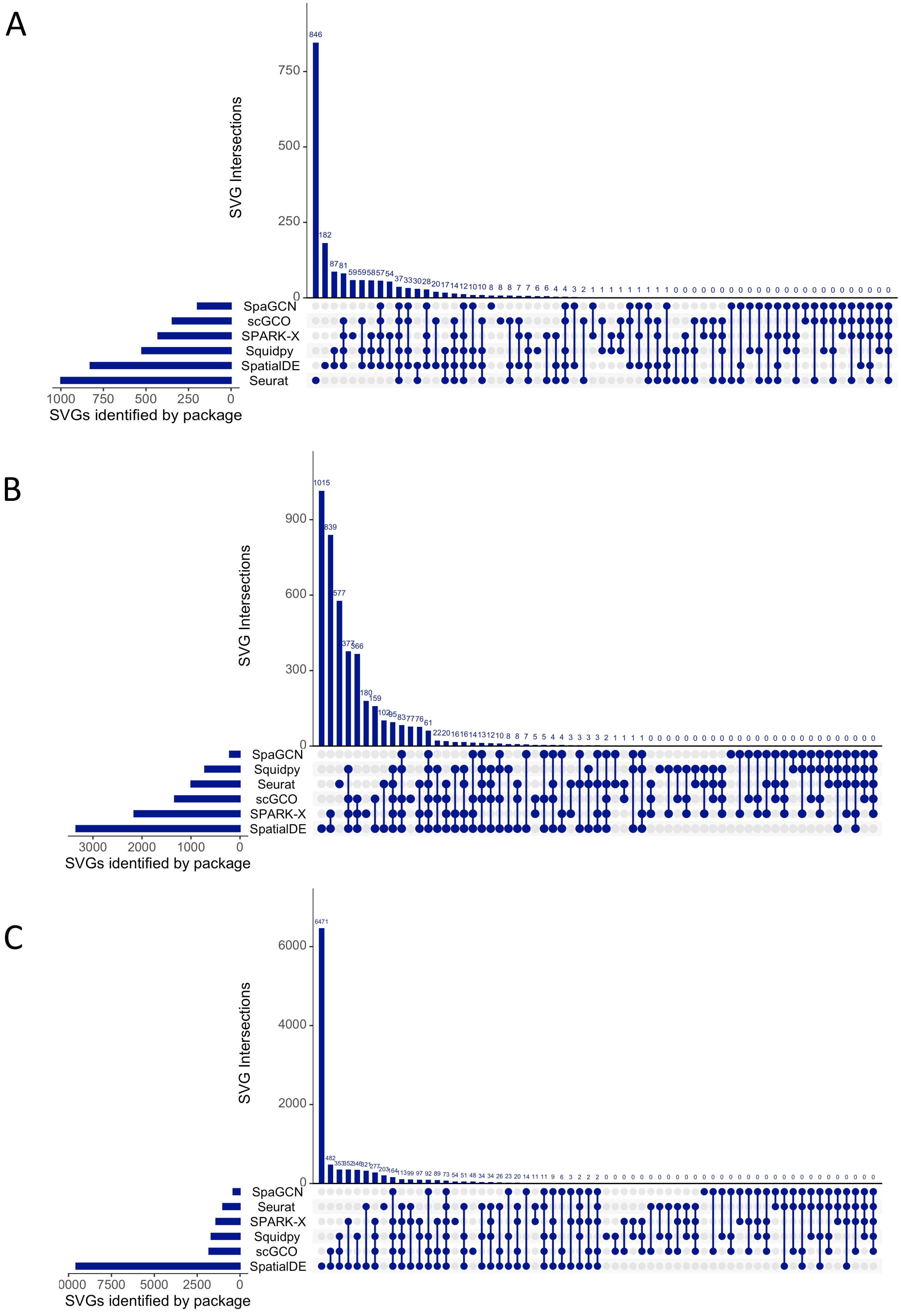

### Additional File 3

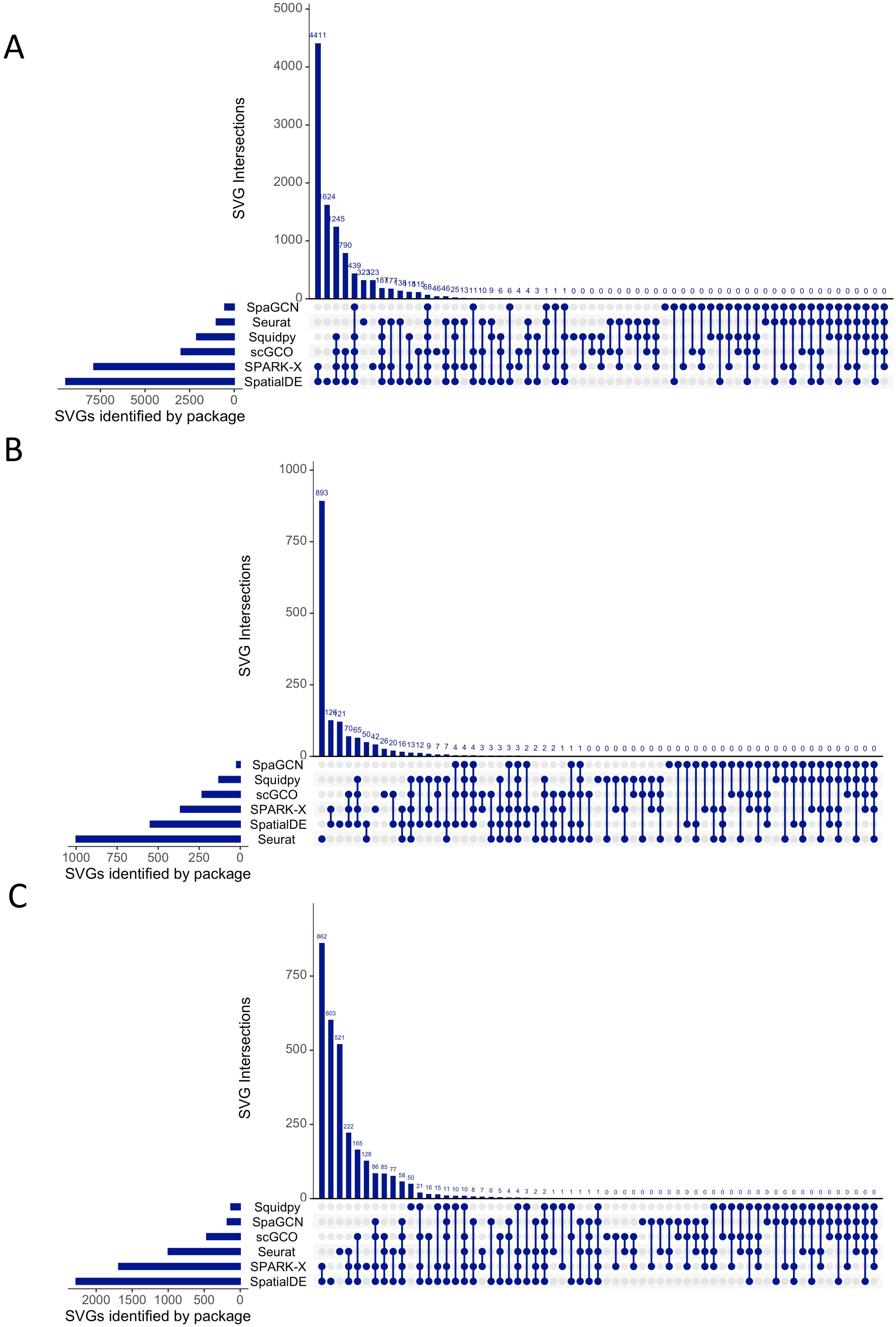

### Additional File 4

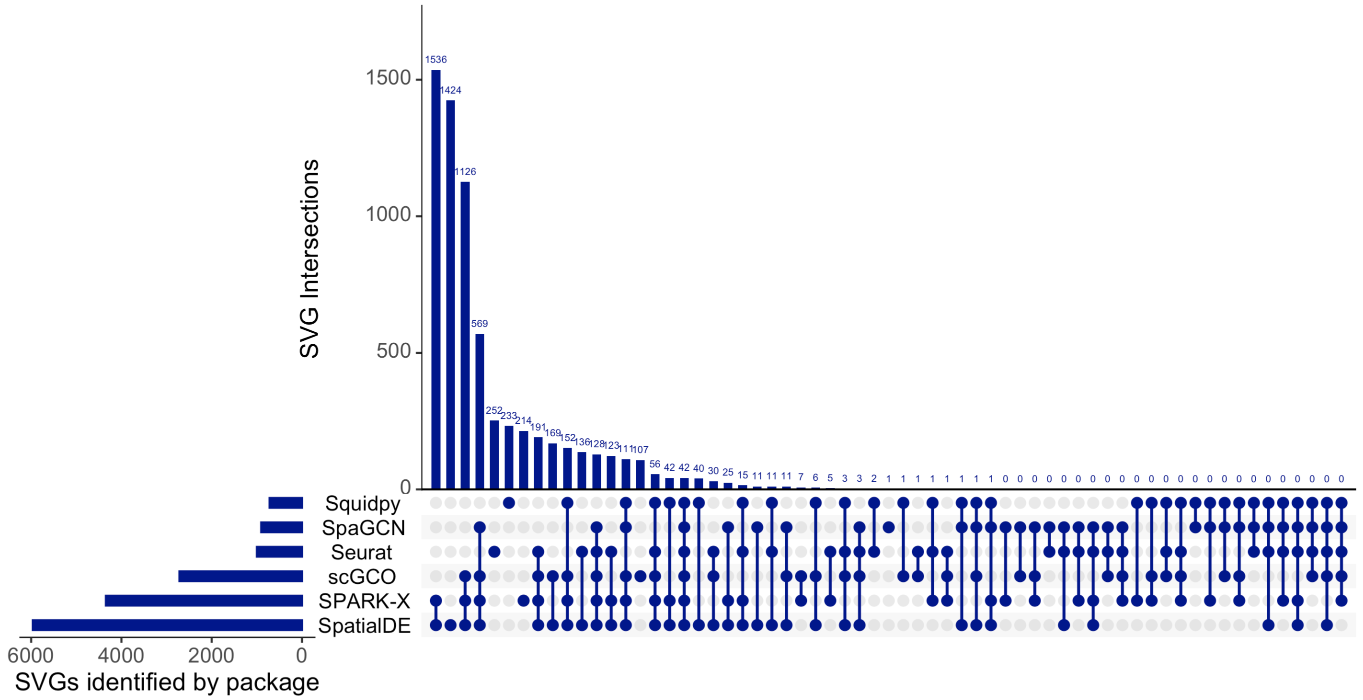

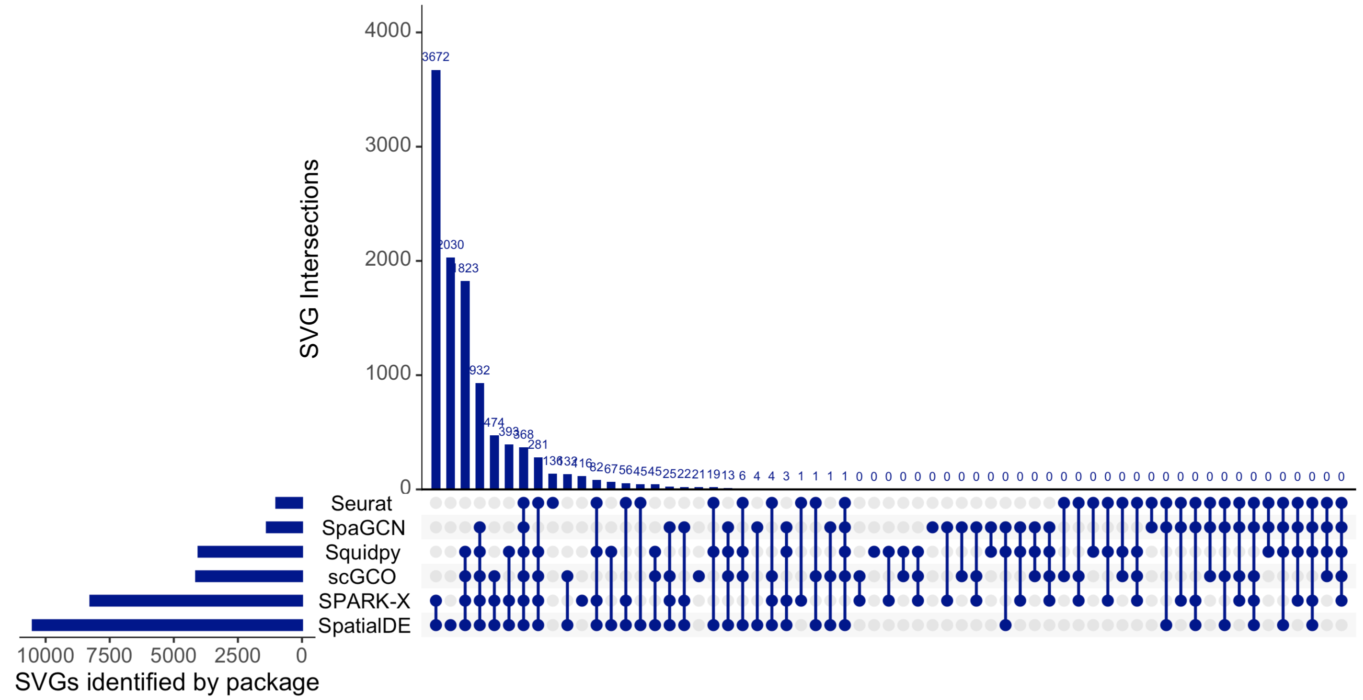

### Additional File 5

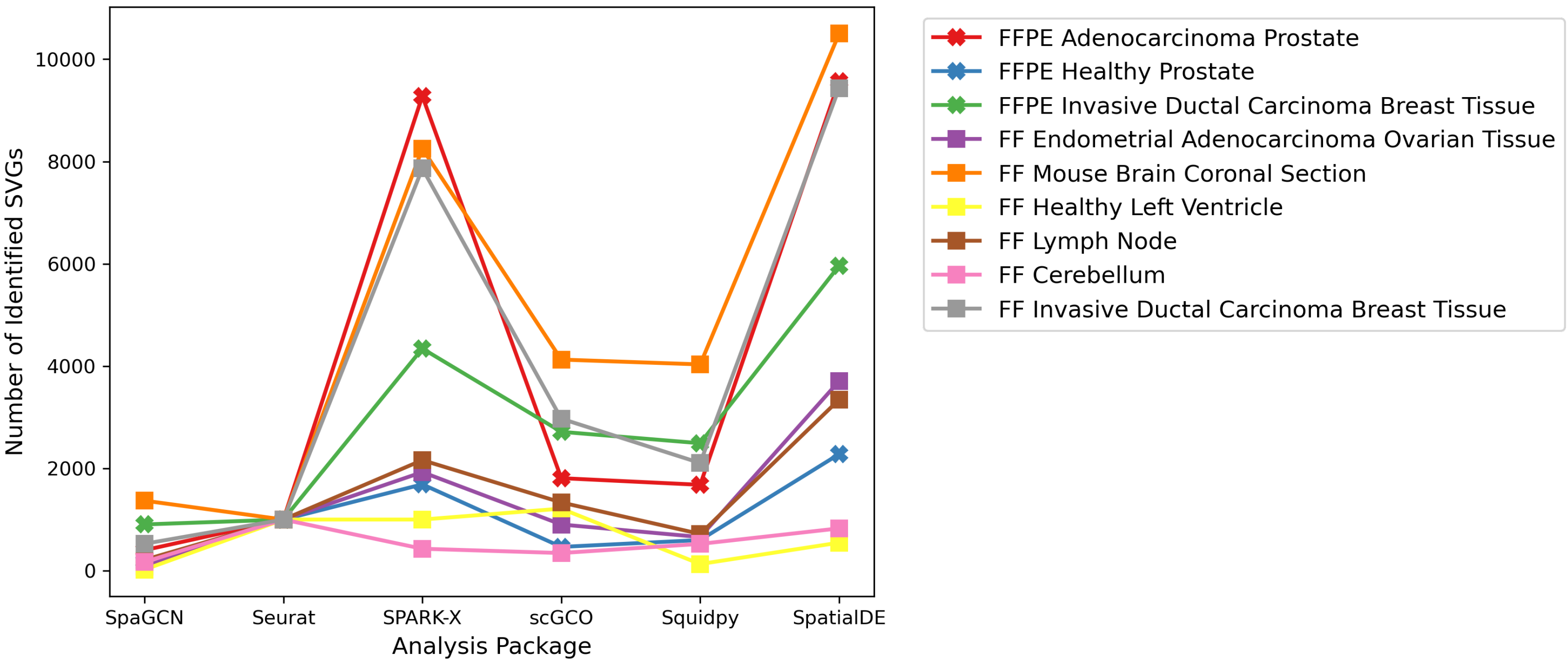

### Additional File 6

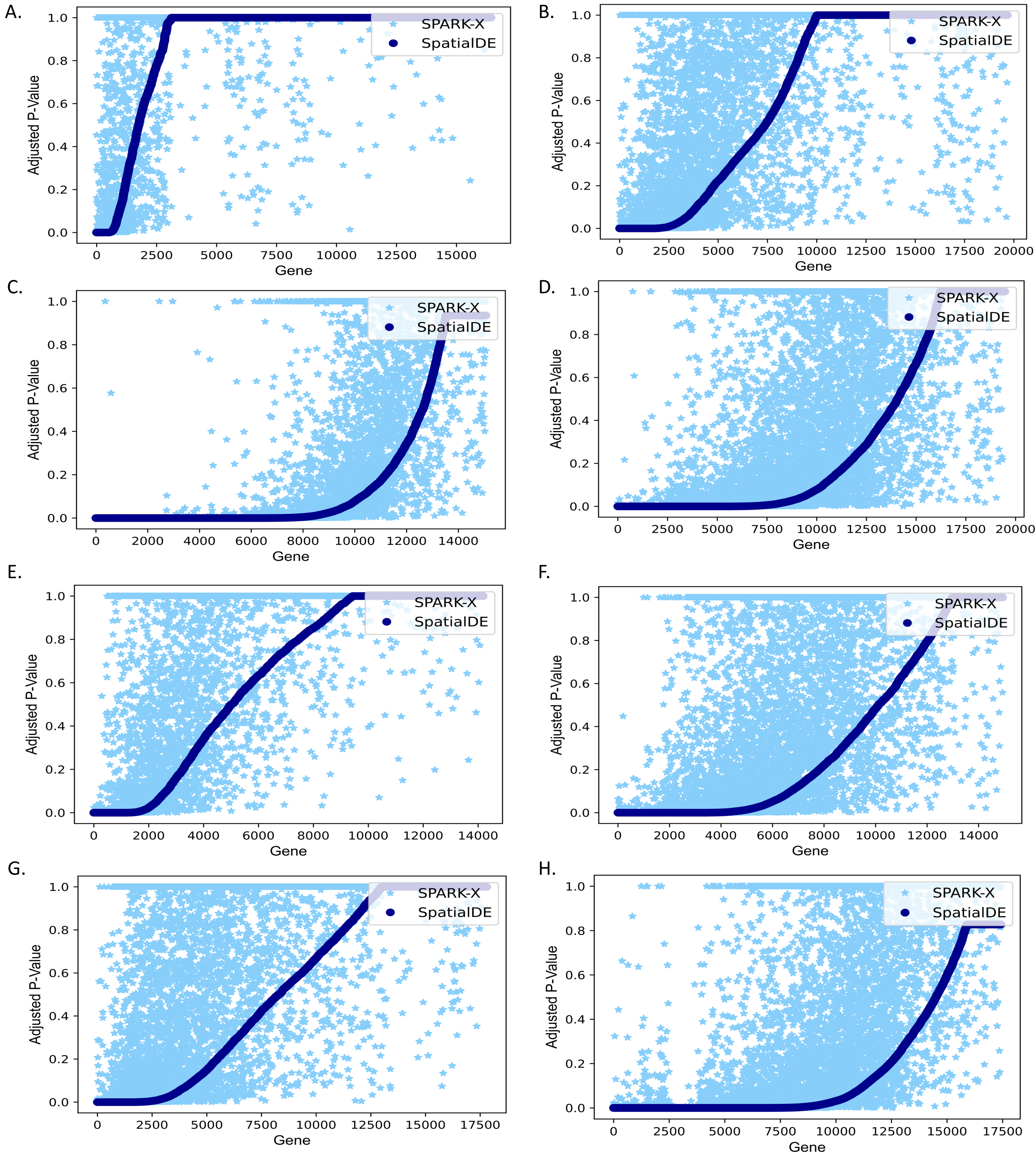

### Additional File 7

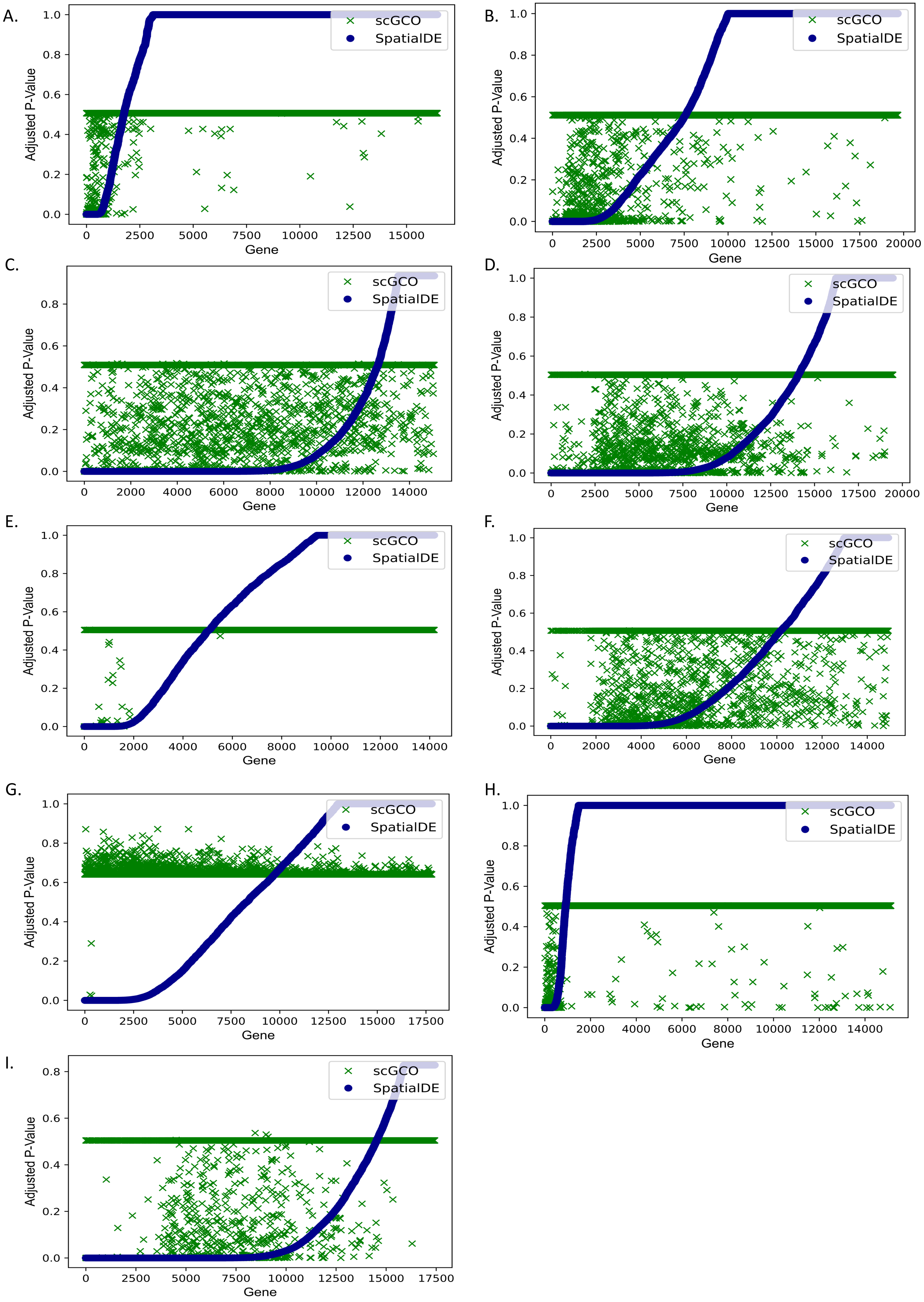

### Additional File 8

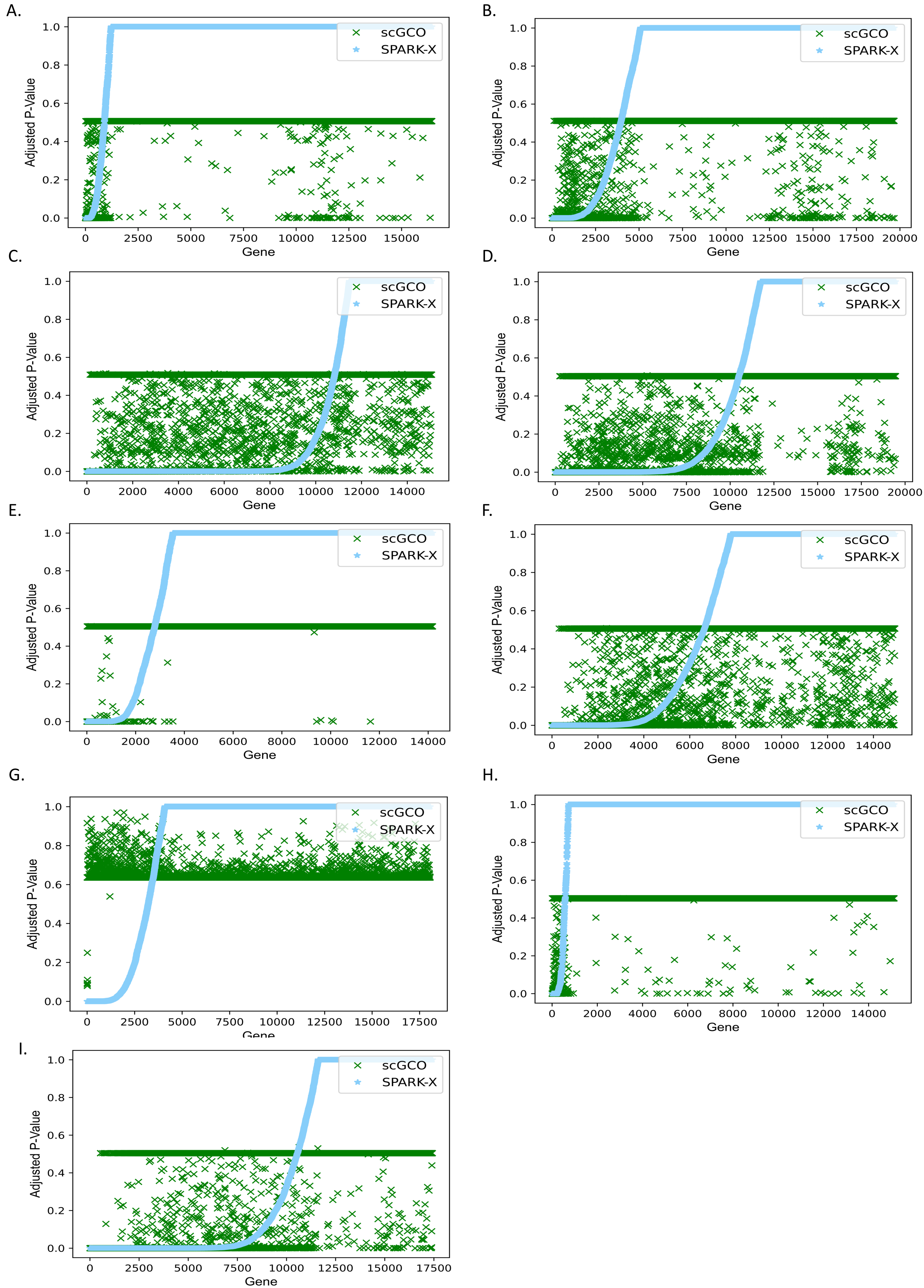

### Additional File 9

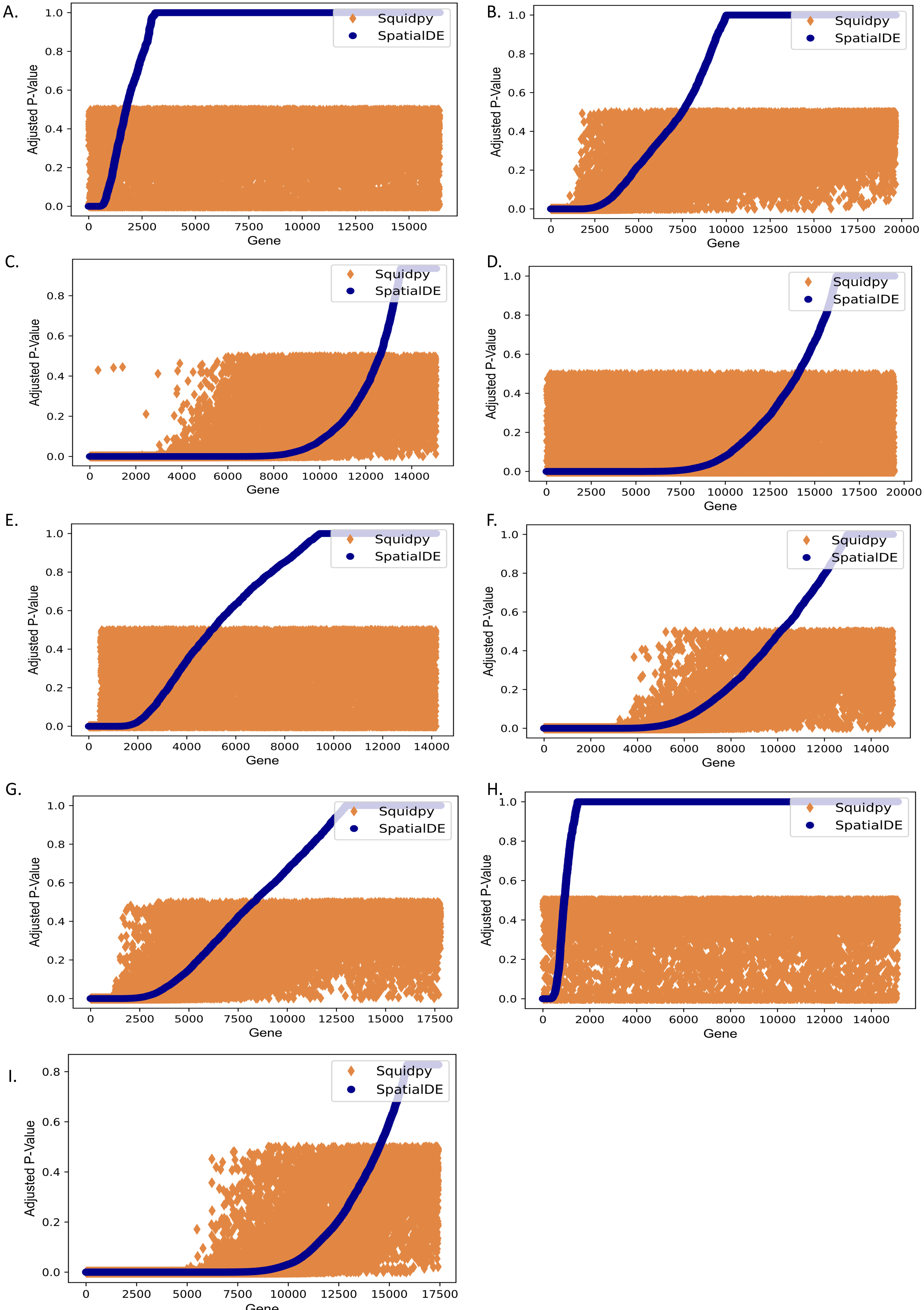

### Additional File 10

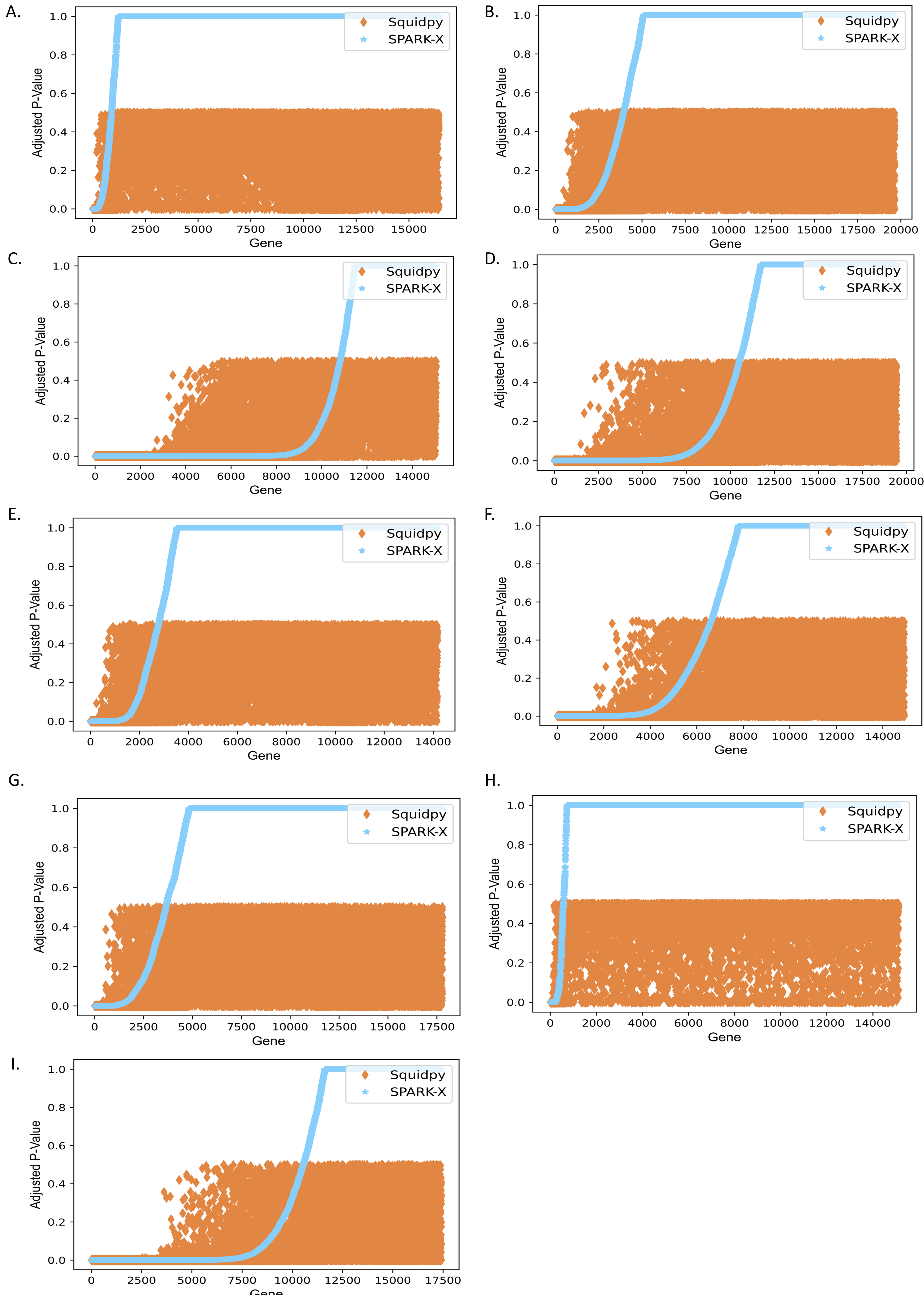

### Additional File 11

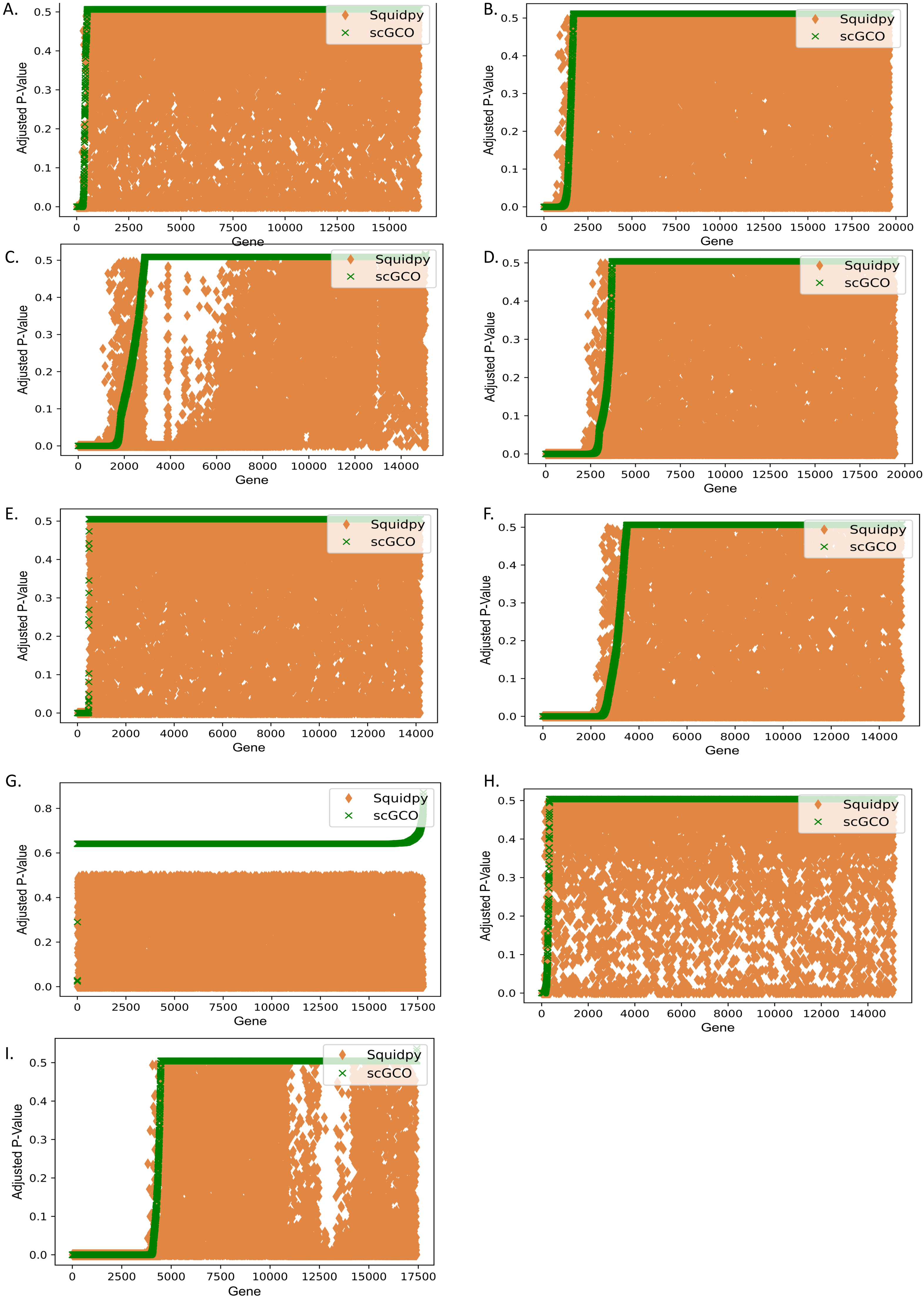

### Additional File 13

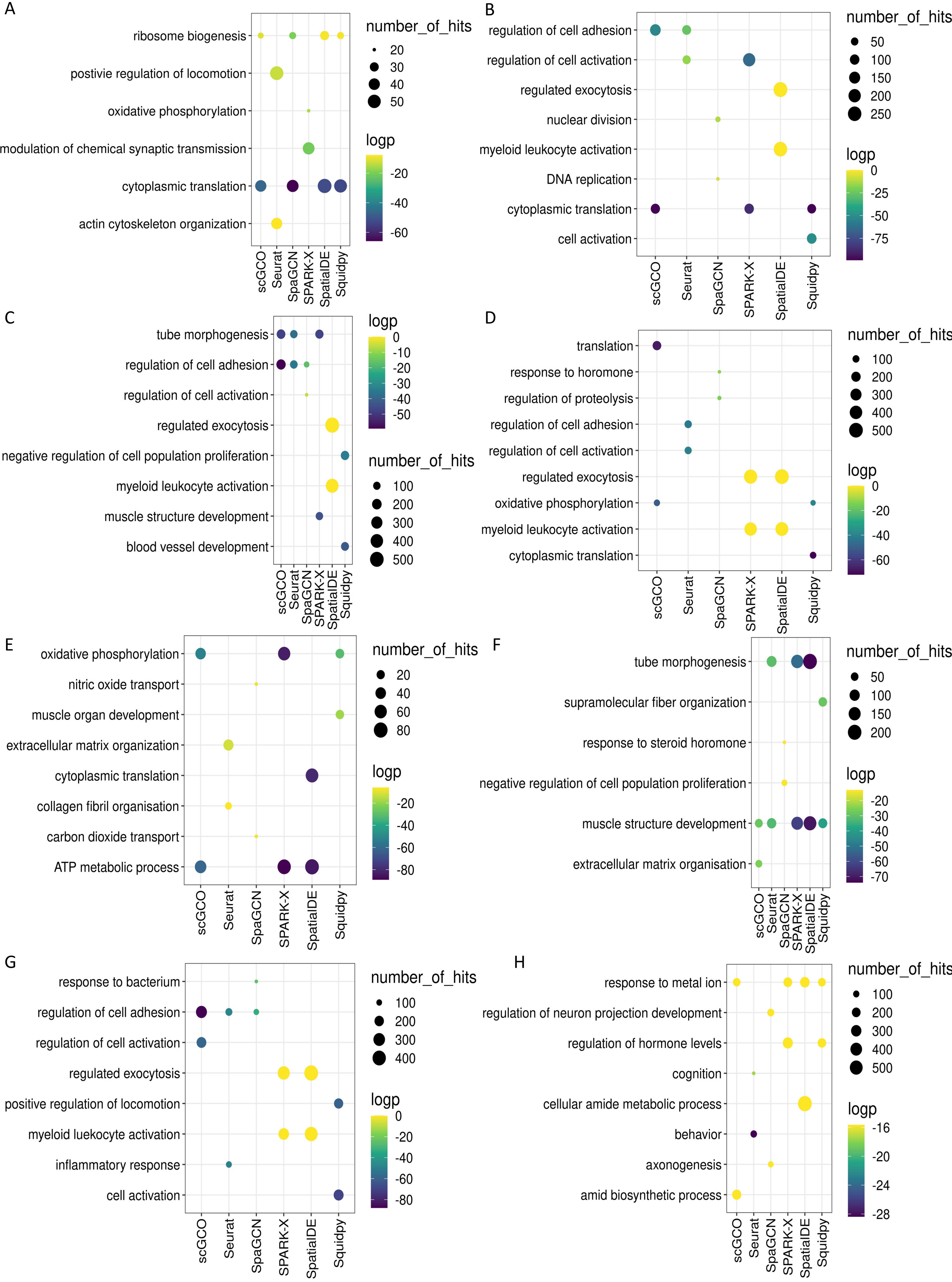

### Additional File 14

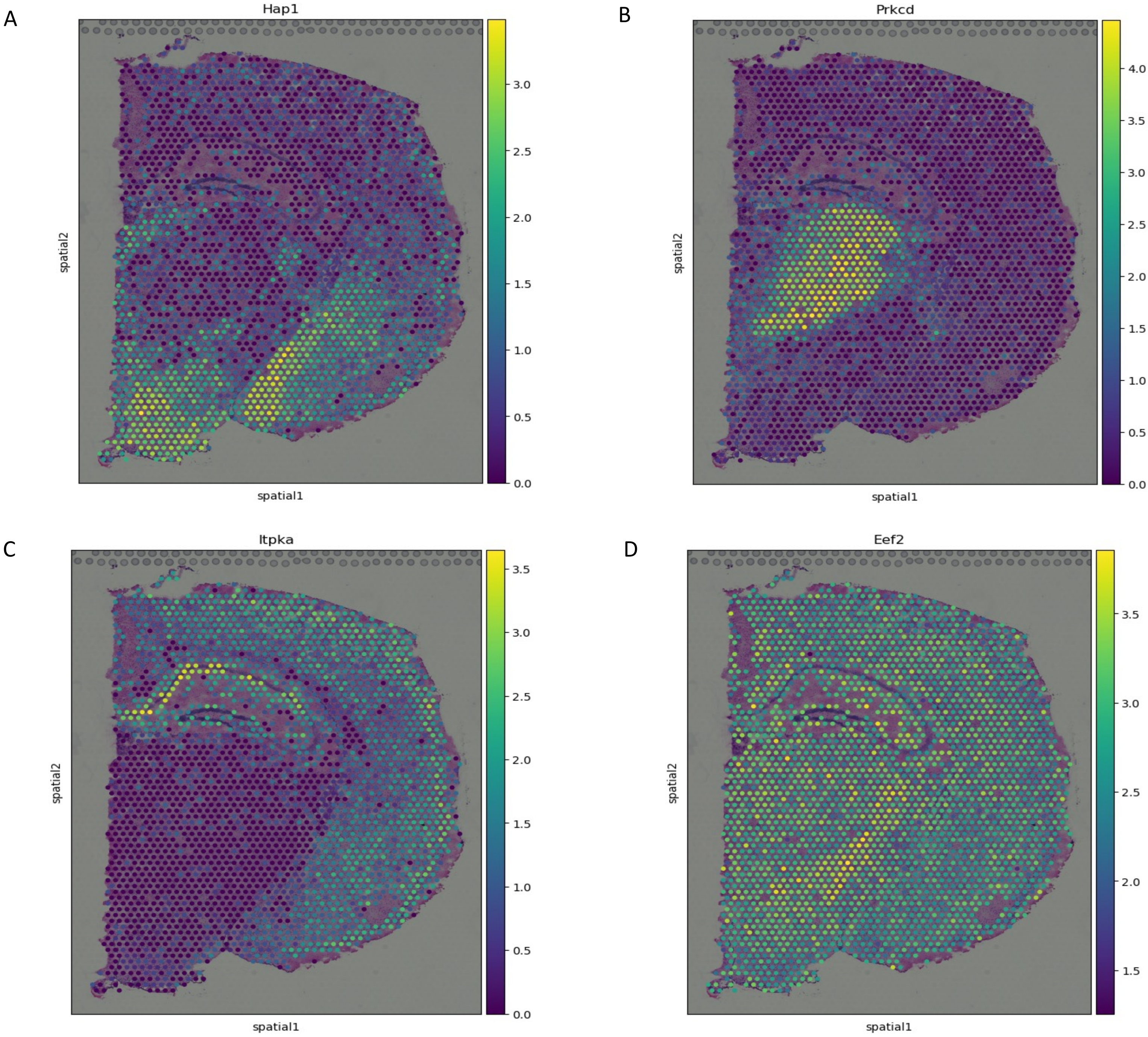

### Additional File 15

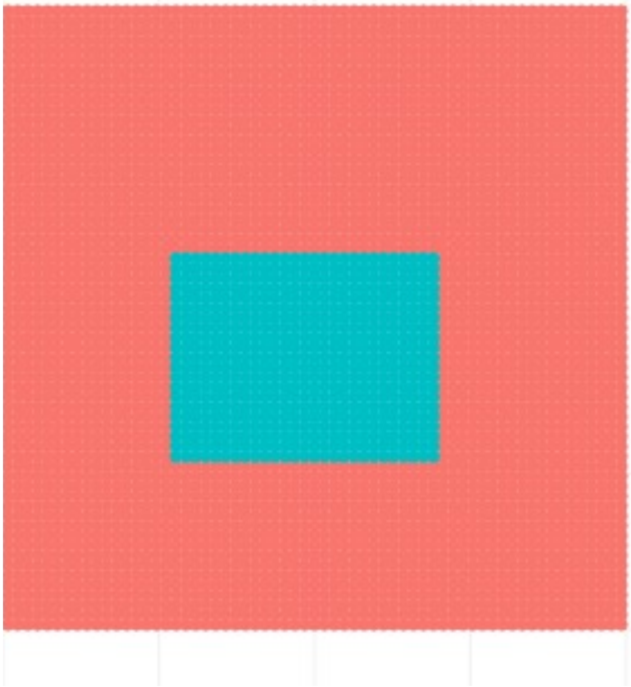

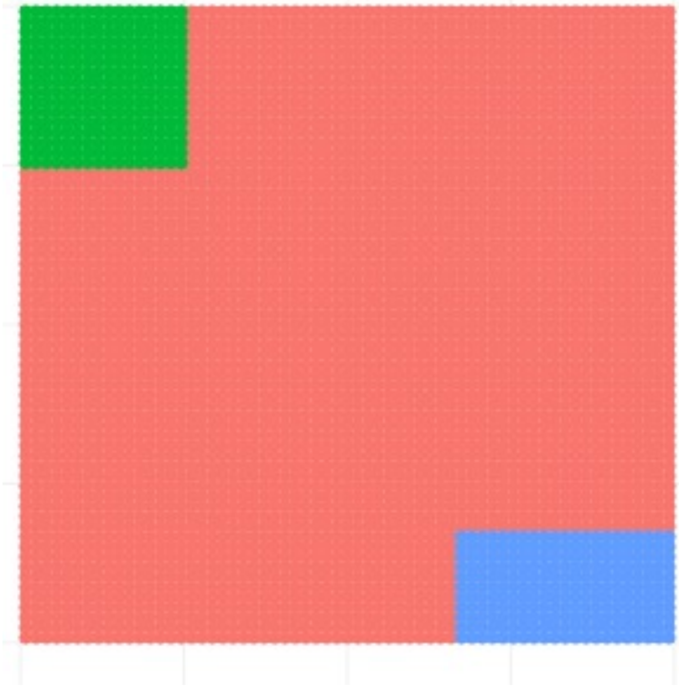

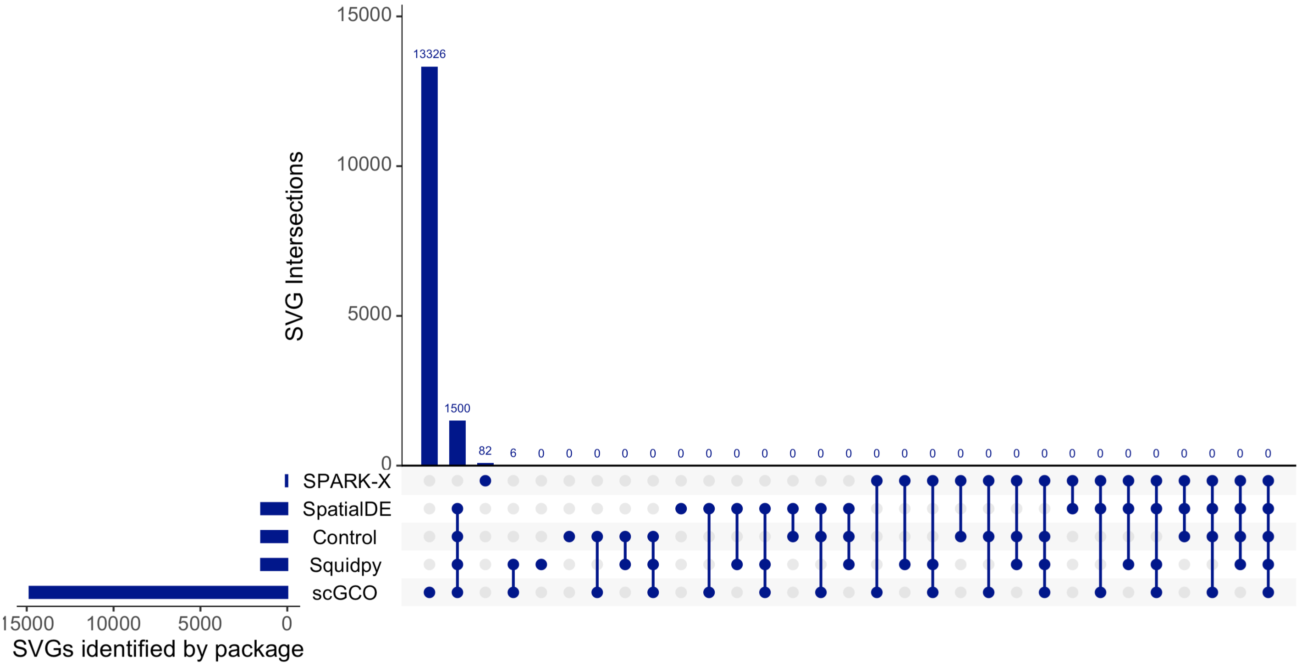

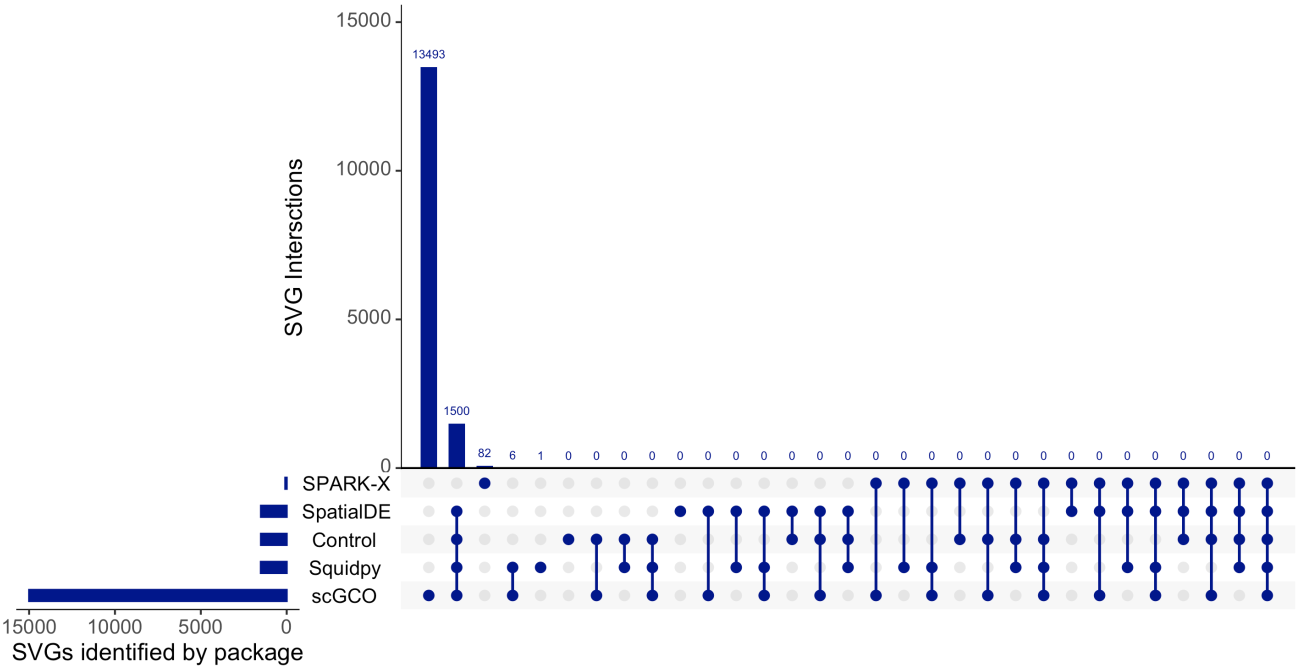

### Additional File 16

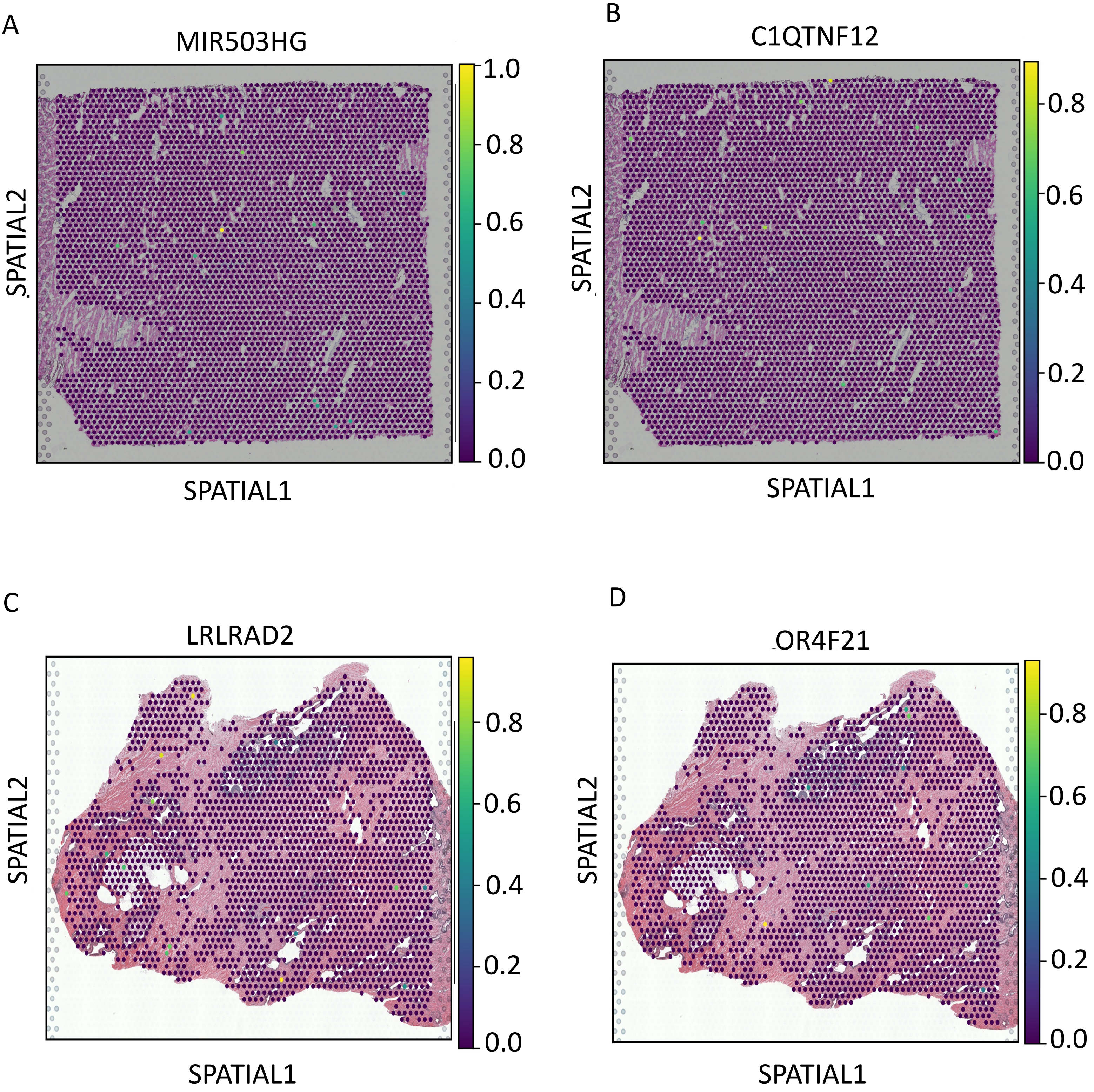

### Additional File 17

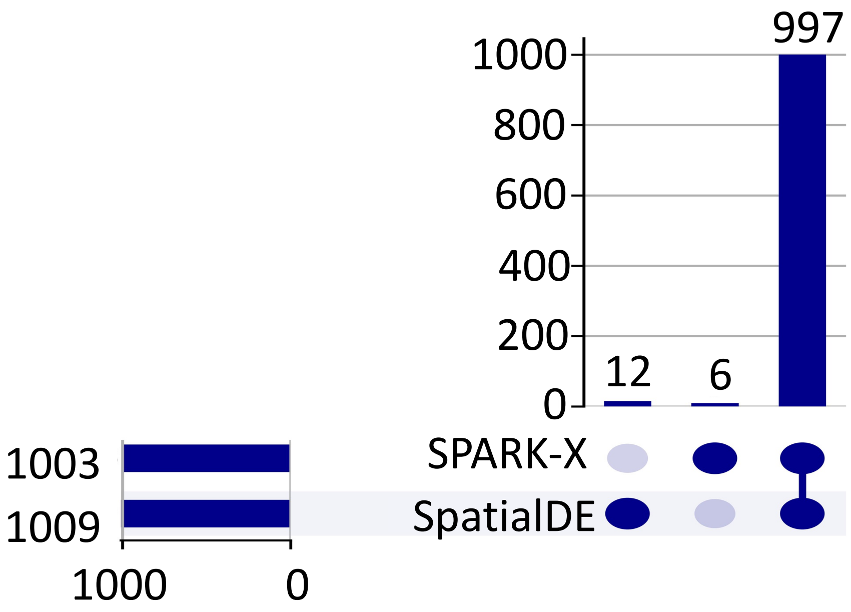
