## Additional File 18 for "Disparities in spatially variable gene calling highlight the need for benchmarking spatial transcriptomics methods"

Supplementary Methods

The aim of this study is to compare publicly available tools that were developed to identify SVGs within spatially resolved transcriptomics (SRT) datasets. When individual tools are published their performance is reported across SRT datasets generated using various platforms, but a systematic comparison of package performance to identify SVGs in FF and FFPE preserved healthy and diseased tissues is lacking. Here, we elected to focus on comparing package performance on data generated using the 10X Visium platform, due to the early commercial availability of the Visium platform which enables the generation of multiple datasets from different tissues and fixation protocols.

Six state of the art packages built for the identification of SVGs (Seurat and Squidpy excepted) in SRT data were selected for benchmarking: SpatialDE, SPARK-X, Seurat, SpaGCN, Squidpy and scGCO. Each package uses a different algorithm to identify SVGs and holds varied assumptions on the distribution of gene expression data (Additional File 12: Table S3). Packages were purposefully chosen to compass a variety of algorithms and associated mathematical models of gene distribution expression that can be used to identify these spatial patterns. While two packages used algorithms based on graph theory, the others were selected to test different mathematical assumptions regarding SRT data. SpatialDE employs Gaussian process regression, a non-parametric probabilistic model (Additional File 12: Table S3) (1). SPARK-X is another non-parametric method, building on a covariance test framework, specifically the projection covariance function (2). This function can measure similarity between two locations or gene expression, it can quantify if the product of the two inputs deviates significantly from the mean value (2). Seurat employs a mark point process, first used by the Trendsceek package (3,4). This is another non-parametric approach that can test if gene expression levels are significantly dependent on the spatial distribution of spots as a function of the distance between them (3). This can then calculate the mark-variogram (3,4). SpaGCN’s method is built around a graph convolutional network (GCN)-based approach and is unique in that it incorporates signal from histology images and restricts identification of SVGs to within spatial domains (5). Gene expression and image data are converted into a weighted undirected graph, which then calculates the distances between any two vertices in the graph (5). The nodes represent spots, and the edge weight is calculated using the histology image and the Euclidian distance between two vertices (5). The weight of each edge is calculated as a function between how related spots are in the graph (5). Dimensionality reduction then graph convolution is computed to then identify spatial domains (5). Finally, differential expression analysis is performed between spots in one spatial domain with neighbouring domains utilising a Wilcoxon rank-sum test and genes with an adjust p-value < 0.05 are reported as SVGs (5). scGCO is another graph-based method, but this package utilises a probabilistic graph model by optimising hidden Markov random fields (HMRF) for the purpose of SVG identification (6). Similar to Seurat, scGCO uses a marked point process to model spatial gene expression (4,6). The dependency of gene expression states on spatial locations is analysed using the complete spatial randomness framework, and scGCO overcomes its limitations using HRMF (6). Finally, Squidpy employs the spatial autocorrelation metrics of Moran’s I to label SVGs (7). Given a continuous feature, in this case gene expression level and their spatial location it can evaluate whether a pattern is present or not using Moran’s I (7).

Each package was tested on nine publicly available, V1 Chemistry Visium datasets generated from healthy and cancerous human tissues, along with a single mouse dataset (Additional File 12: Table S4). Additional testing was performed using simulated data (Additional File 1). The filtered output files and imaging data from the Space Ranger v1.0.0 pipeline were downloaded for each dataset. Further filtering and pre-processing of the data was performed using Scanpy v1.8.1 and the following were removed: genes expressed in fewer than 10 spots, spots with fewer than 2000 counts, and/or spots with fewer than 2000 genes expressed. As the percentage of mitochondrial gene expression and maximum number of counts per spot varied significantly between datasets, filtering was performed on a per-dataset basis. Mitochondrial filtering in FF Cerebellum was limited to spots with less than 15% mitochondrial genes (8). Counts were then normalised per spot, log transformed, and the top 2000 highly variable genes identified. This count matrix was used as the input files for each algorithm. A custom script was written for conversion between the Scanpy AnnData object and the Seurat S4 object input.

Analysis using SpatialDE was performed using default parameters using Python v 3.9.7. Reported SVGs were filtered to include only those with an q-value <= 0.05.

Analysis using SPARK-X v 1.1.1 was performed with default parameters using R v 3.6.1. Reported SVGs were filtered to include only those with an adjusted p-value <= 0.05.

Analysis using Seurat v 4.1.0 was performed with default parameters using R v 3.6.1. A custom script was used to convert the Anndata to Seurat object. SVGs were identified using the function FindSpatiallyVariableFeatures with default parameters as specified in the vignette (top 1000 variable features selected and markvariogram selection method).

Analysis using SpaGCN v 1.2.0 using default parameters after calculating the appropriate radius for each dataset using Python v 3.9.7. Number of target domains was informed by Scanpy clustering and SVGs that were common between domains were removed.

Analysis using scGCO v 1.1.1 with default parameters using Python v 3.7.6. Reported SVGs were filtered to include only those with a FDR value <= 0.05.

Comparison of the number of SVGs identified by each package across all datasets visualised in Fig. S5 (Additional File 6), were modified from code generated by ChatGPT (9).

Upset plots were generated using package upsetplot v 0.6.0 and Python v 3.9.7. Pattern of SVG expression were generated using Scanpy v1.8.1. The upset plots display the unique intersection of different overlapping or independent results identified across packages within a single dataset and serve to highlight the differences in the scale of the number of SVGs identified within a dataset. Furthermore, visualisation of datasets with the most reported SVGs independent of the other packages may highlight the introduction of FP.

The SVGs identified by each package were used as the inputs for gene ontology enrichment analysis. When less than 3000 SVGs were identified, Metascape was used for analysis, while in instances where there were over 3000 inputs WebGestalt was used as Metascape has a limit on the number of genes accepted as inputs. Metascape was run with express analysis and the input as species was set to *H.sapiens or M.musculus*. For WebGestalt analysis the organism of interest was set as *Homo sapiens* or *Mus musculus*, method of interest as over-representation analysis, functional database of ‘gene ontology’ focusing on biological process and the ‘reference set’ was set as ‘genome’. All results were downloaded then filtered down to the top 10 significant GO terms. ggplot2 package was used to visualise GO terms.

Seurat ranks highly variable genes by how dependent their expression is on spatial location, then produces a list of SVGs (4). SpaGCN reports unraked lists of SVGs with a p-value of < 0.05. For the four packages that generated p-values associated with the identified SVGs (SpatialDE, Squipdy, SPARK-X and scGCO) a Wilcoxon signed rank test was performed. For each of the nine publicly available datasets, each possible pairwise combination of the results from different packages was analysed using the scipy stats function to perform a Wilcoxon signed rank test. The concordance for each dataset was calculated using a pairwise comparison between the results of SpatialDE, Squidpy, SPARK-X and scGCO for all available combinations.

Simulated Datasets

Simulated datasets with known patterns of SVGs were generated using SRTsim (10). SRTsim allows the user to generate reference-free, simulated data that captures many of the features typical of Visium datasets through a Shiny app interface (10). These features include the number of spots and genes in the dataset, the mean and dispersion values of gene expression as well as the proportion of zero counts (10). If regions are selected within the dataset, the log fold-change within this region can be set compared to the rest of simulated area. Expression counts were simulated using a negative binomial distribution for both datasets (10). Two datasets were generated using SRTsim for this purpose. A square grid is selected (reproducible seed 2), with 4,000 spots and 15,000 genes overall. A square ‘hotspot’ pattern of 675 spots was generated in the centre of the grid, which will serve the area of higher gene expression for the simulated SVGs (Additional File 15). 1,500 signal genes were then simulated with 13,500 noise genes, no lower signal genes were selected. A reproducible seed of 2 was set and baseline overdispersion was set to 0.5 with the baseline mean was set to 3. A second simulated dataset was generated with SRTsim with a slightly more complex gene expression pattern. The grid layout and number of spots was the same as the aforementioned dataset, set with reproducible seed 3. Here two separate regions were generated to contain simulated SVG expression, in opposite corners of the square layout (Additional File 15). The two groups had a log fold-change of 5 compared to the rest of the area (Group A) with a log fold-change of 1. Next the expression of 15,000 genes were simulated across the dataset: 13,500 noise genes, 750 high signal SVGs and 750 low signal SVGs. Dispersion and mean values were the same as the previous dataset. This second dataset was generated with the aim of showcasing which packages may be adept at identifying SVGs with a lower signal across the dataset or a pattern different to the typical ‘hotspot’ which is often shown, such as in early simulations available with SPARK-X (2). The overlap of the results from each package with the known SVGs and noise genes in the data was then calculated to determine the sensitivity and specificity of each package. R scripts to calculate the sensitivity and specificity of each package were modified from code generated by ChatGPT (9).

A total of four negative control datasets were generated, two by randomising the values of the FF left ventricle data and two by randomising the values of the FFPE prostate data in python. For each of the two datasets, one simulated dataset was created by randomising the x and y spot coordinates for this dataset independently to remove spatial correlation. Subsequently, an additional simulated dataset was generated from each dataset by randomising the spatial coordinates and each column in the gene expression matrix to ensure any overall associations were removed. To generate a simulated dataset of known coordinates, the code available in the SPARK-X simulated data vignette was used. To test all simulated datasets the same parameters for each algorithm were used as in the analysis of the publicly available datasets. The true positives rates and false positive rates were then calculated for the results all packages that were compatible with the dataset.
